# Ubiquitin-mediated proteolysis in *Xenopus* extract

**DOI:** 10.1101/062331

**Authors:** Gary S. McDowell, Anna Philpott

## Abstract

The small protein modifier, ubiquitin, can be covalently attached to proteins in the process of ubiquitylation, resulting in a variety of functional outcomes. In particular, the most commonly-associated and well-studied fate for proteins modified with ubiquitin is their ultimate destruction: degradation by the 26S proteasome via the ubiquitin-proteasome system, or digestion in lysosomes by proteolytic enzymes. From the earliest days of ubiquitylation research, a reliable and versatile “cell-in-a-test-tube” system has been employed in the form of cytoplasmic extracts from the eggs and embryos of the frog *Xenopus laevis*. Biochemical studies of ubiquitin and protein degradation using this system have led to significant advances particularly in the study of ubiquitin-mediated proteolysis, while the versatility of *Xenopus* as a developmental model has allowed investigation of the *in vivo* consequences of ubiquitylation. Here we describe the use and history of *Xenopus* extract in the study of ubiquitin-mediated protein degradation, and highlight the versatility of this system that has been exploited to uncover mechanisms and consequences of ubiquitylation and proteolysis.

## Introduction

Proteins are created through transcription and translation from DNA to RNA, and then finally synthesized at the ribosome [1]. In addition to their production, proteins are also destroyed to regulate their levels, terminate their function and recycle their constituent amino acids for future protein syntheses. The study of protein catabolism is a relatively recent area of research, as despite early observations that proteins were broken down and their components reused [2, 3], there was significant resistance to the concept that proteins were not persistent, and were degraded intracellularly [4]. It was only after extensive investigation [5, 6, 7] that protein degradation came to be accepted, and the ubiquitin (Ub) protein was identified as a mediator of ATP-dependent protein degradation [8, 9].

Ub can be covalently attached to other proteins in the post-translational modification known as ubiquitylation. Ubiquitylation of proteins can result in a variety of signaling roles [10], but the best studied outcome of ubiquitylation is protein degradation. Ub-mediated proteolysis can be achieved through degradation by the 26S proteasome through the Ub-Proteasome System (UPS) [11], or by digestion in organelles called lysosomes using proteolytic enzymes [12].

The study of protein degradation and ubiquitylation continues to be a highly active and continuously-evolving field (as reviewed in [13]), one which has taken advantage of a number of model systems, and for biochemical studies, the frog *Xenopus* has played an important role. In particular, *Xenopus* offers the unique advantage of the ability to generate cell-free cytoplasmic extracts, which contain soluble proteins capable of carrying out the biochemical modifications required for protein ubiquitylation and destruction, facilitating many important advances in this field. Here, we describe the *Xenopus* cell-free cytosolic extract system and illustrate its utility in the study of protein ubiquitylation and degradation.

## *Xenopus* extract systems

*Xenopus* is an ideal model organism for biochemical studies due to its unique tractable extract system; milliliter volumes of cytosol readily generated from the thousands of eggs a single female frog can lay in one day can be used to study ubiquitylation and protein destruction events *in vitro*. Cytosolic extracts were first generated using unfertilized eggs from *Xenopus laevis* to demonstrate the assembly of chromatin from DNA [14], but have been readily adapted for study of a variety of biochemical and cell biological regulatory mechanisms. The extracts that are used most commonly to investigate mechanisms of ubiquitylation are “cell cycle” extracts. These have been further modified over time to suit the requirements of the researcher, and often developed from the protocols laid down by Andrew Murray [15].

Activated interphase cell cycle extracts, made using the general protocol illustrated in (Figure 1), were first developed by Manfred Lohka in the lab of Yoshio Masui using *Rana pipiens* cytosol to study pronuclear formation with *Xenopus laevis* sperm nuclei [16, 17]. Modified versions, which still retained activity after extract was frozen and thawed, were used to study DNA replication [18] and, as we will discuss later, to study cyclin degradation. These studies involved both “low-speed” extracts (generated after low speed centrifugation that including light membranes, ribosomes and nuclear envelope [19]) and “high-speed” extracts (generated after an additional second, faster centrifugation to remove membranes and ribosomes [20]). Extracts that could go through multiple cell cycles were also developed [21, 22]. In addition, non-activated or cytostatic factor (CSF)-arrested meiotic metaphase extracts have been generated, replicating the environment of the mitosis-like state of unfertilized egg and these could later be activated and driven into interphase with the addition of calcium ([23], with further refinements [24]). Protocols for both activated interphase and CSF-arrested mitotic extracts have been described in detail by Murray [15]. In addition to egg-based extracts, we have also used embryonic extracts, generated by centrifugation of embryos at particular stages of development, to mimic developmental stages such as neurogenesis [25]. Protocols for the preparation of a variety of cytosolic extracts to answer many experimental questions can be found at the community portal for the *Xenopus* community, Xenbase (www.xenbase.org/, [26, 27]).

The utility of the *Xenopus* extract system to study a variety of biological problems is clear (e.g., [15, 28, 29] and many others). Here we are focusing on its use in the study of proteolysis, where the simplicity of the ubiquitylation and degradation assays that the extract system allows has been key to the significant advances that have resulted (see [30, 31, 32] for examples of Ub protocols). In addition, it has been straightforward to determine how biochemical mechanisms relate to the situation *in vivo* by comparing with the developmentally tractable *Xenopus* embryo system. Importantly, mechanisms identified in frog have been widely shown to be conserved in mammalian systems, demonstrating the utility of this “cell-in-a-test-tube” analytical approach.

## The machinery of ubiquitylation

The pathway leading to UPS-mediated protein degradation is now well characterized and is discussed in more detail elsewhere ([33, 34, 13, 35]. In brief, Ub is activated by an E1 enzyme [36], passed to an Ub-conjugating (E2) enzyme [37] and from there to a substrate protein, specified by the particular E3 ligase that binds both the substrate and the E2 [38]. Multiple rounds of ubiquitylation on Ub itself can then follow to create chains of polyUb. The polyUb modification most commonly understood to target proteins for UPS-mediated degradation is a tetramer of K48-linked Ub moieties [39]. However many novel chains, of homotypic, heterotypic, and even branched topologies are contributing to the study of “atypical” chain ubiquitylation [40]. It was observed that a purified cell-cycle dependent component of the ubiquitylation machinery, the Anaphase-Promoting Complex/Cyclosome (APC/C) (see below), from *Xenopus* extract at different salt conditions, could bring about the formation of Ub chains of differing lengths. In particular it was demonstrated *in vitro* at least, that the APC/C could generate K11-, K48- and K63-linked polyUb chains targeting protein substrates for degradation [41]. Upon further refinement, this led to work on the precise mechanism for the formation of K11-linked chains by the APC/C using the specificity conferred by the E2, UBE2S [42]. This and other work demonstrates that the *Xenopus* system has played a central role in identifying which proteins are targeted for degradation and when.

**Figure 1:**
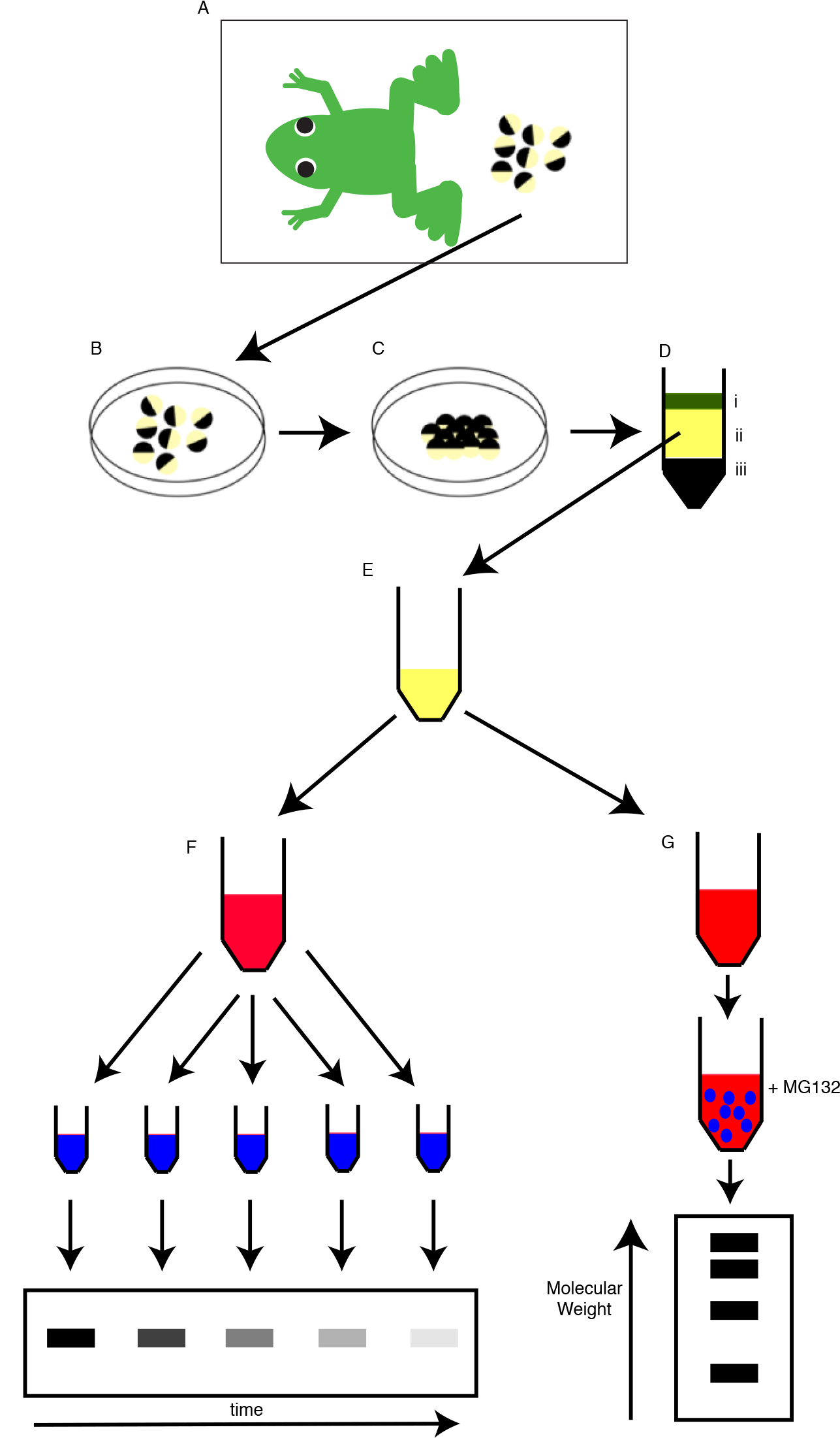
*Xenopus laevis* egg extract systems in ubiquitylation studies. (A) Female *Xenopus laevis* are induced with hormone to lay eggs; (B) eggs are collected and their jelly coats removed; (C) (optional) eggs are activated using calcium ionophore to mimic fertilization and direct entry into interphase. Activated eggs are then spun at high speeds in a centrifuge to separate into (D) (i) lipid, (ii) cytoplasmic, and (iii) pigment granule and yolk platelet layers. (E) The cytoplasmic layer is taken and used for *in vitro* studies: for example, (F) *in vitro* translated radiolabelled protein is added to extract supplemented with Ub and ATP, and aliquots removed at various time points to assay degradation of protein over time; and (G) *in vitro* translated radiolabelled protein is added to extract supplemented with tagged Ub and the proteasome inhibitor MG132, and after an incubation period antibody-coated beads are added to isolate ubiquitylated proteins under various conditions.

## Discoveries of Degrons and cell cycle regulated proteolysis

The use of *Xenopus* extracts has been central to our understanding of mechanisms that regulate the destruction of proteins to bring about cell cycle transitions; they can readily be manipulated to mimic the cellular conditions of interphase and mitosis and it has been known for a long time that transition between these cell cycle states is accompanied by the cyclical synthesis and destruction of a number of proteins [43].

Global Ub metabolism is not obviously regulated by the cell cycle; using cycling *Xenopus* egg extract, it was possible to show that the levels of Ub did not change with the cell cycle phase; the rates of conjugation, protein degradation, and isopeptidase activity all remained constant throughout the course of the cycling extract experiments [44]. However, some proteins such as cyclins are destroyed at specific points in the cell cycle [45]. The authors concluded that, therefore, cell cycle differences in cyclin degradation could not be accounted for by any obvious global differential Ub kinetics in different phases of the cell cycle, but instead must result from individual regulation of specific proteins. In fact, the experiments by Glotzer *et al.* suggested that the recognition of cyclin B by the ubiquitylation machinery itself promotes the entry of the cell cycle into anaphase [45]. (Prior to this work, only one other case of physiological regulation through Ub-mediated degradation had been identified: the plant protein phytochrome [46]). In their landmark study, Glotzer and colleagues found that the N-terminal region of cyclin B was required for its degradation in a mitotic extract system and that cyclin B was itself ubiquitylated [45]. In the absence of inhibitors of ubiquitylation or proteasomal inhibitors (not yet described at that time), the authors were able to demonstrate the link between ubiquitylation and degradation of cyclin B by calculating the kinetics of protein degradation compared with the flow of cyclin B through ubiquitylated intermediates [45].

As well as demonstrating the role of cyclin B destruction in the cell cycle, these and further *Xenopus* extract experiments have shown us that cell cycle regulated destruction of proteins such as cyclins relies on timed degradation orchestrated by sequences known as degrons [45, 47, 48, 49, 50]. Degrons are the minimal signals within proteins that result in their targeting for degradation by the proteolytic machinery [51]. Here we will discuss two well-known degrons identified using *Xenopus* extract systems that are targeted by the E3 ubiquitin ligase Anaphase Promoting Complex/Cyclosome (APC/C see below): the KEN (Lys-Glu-Asn) box and Destruction (D-box, Arg-X-X-Leu).

The D-box was first identified during the early investigations of the degradation of the cell cycle regulator cyclin B, showing that a particular region of the N-terminus of the cyclin was necessary and sufficient to target it for degradation in *Xenopus* mitotic extract; stable entry of extract into mitosis could be brought about by addition of a form of cyclin B with the first 90 amino acid residues deleted (a technique still used to generate mitotic egg extract), while adding this N-terminal portion of cyclin B to other proteins was enough to trigger their proteolytic degradation [45]. The characterization of the precise degron, the D box, within the Cyclin B N-terminal region was further refined using a mutagenesis approach, followed by investigation of the kinetics of protein degradation in *Xenopus* extract systems [47]. Furthermore, important differences between control of cyclins A and B destruction was also identified using egg extracts, resulting from differing residues within their respective degrons [47].

The discovery of the KEN box, a distinct degron targeting proteins for cell cycle regulated destruction in *Xenopus* extracts, highlighted an important mechanism for cell-cycle regulation of protein degradation by the APC/C multi-component E3 ligase (see below). The association of Cdc20 with core components results in the mitotically active form of the APC/C, while the association of Cdh1 generates the interphase form [52, 53, 54]. While Cdc20 mitotic substrates contain the D-box [45], Cdh1 could target both D-box and non-D-box-containing proteins for degradation. As Cdc20 itself lacks a D-box but is degraded in a Cdh1-dependent manner, it was an ideal candidate for identification of other potential degrons, and this resulted in the discovery of the KEN box [48]. Additional substrates such as proteins Nek2 and B99 were also found to be ubiquitylated and degraded by the natural presence of KEN boxes and, like the D-box [45, 47], the KEN box can target heterologous proteins for degradation when fused to them [48].

The anaphase-promoting complex (or APC) was discovered using *Xenopus* extract systems [55] at a similar time to the discovery of the cyclosome in clam oocyte extracts [56]. Now collectively known as the APC/C, the identification of this multi-protein ubiquitylation complex that shows cell cycle stage-specific activation by Cdc20 and Cdh1 (see above), and the identification of cell cycle regulated degrons, combine to explain the cell cycle regulation of cyclin stability. It is not possible to overemphasize the role of *Xenopus* extract in facilitating the discovery of the APC/C and for further elucidating its function; for instance the identification of key binding regulatory proteins such as Cdc20 was made possible using exploratory experiments in *Xenopus* extract along with work in HeLa cell extracts [52]. Similarly, the identification of Mad2L2 as a regulator of Cdh1 dissociation, and the proposal of a mechanism for Mad2-mediated Cdc20 dissociation and Mad2L2-mediated Cdh1 dissociation from the APC/C, were made possible through *Xenopus* extract-based assays of APC/C activity [57]. Identification of subunits such as BIME [58] and additional components such as Fizzy required in APC/C activation [59] were also all absolutely dependent on using *Xenopus* extract systems. It was also used to characterize the kinetics of cyclin ubiquitylation to reveal more general properties of cyclin B protein proteasomal degradation, as described above [45]. For instance, using *Xenopus* extract, Yu *et al.* purified the E2 enzyme UBCx from interphase egg extracts [60], and were thus able to identify and characterize a previously observed Ub-conjugating activity targeting cyclin B [55].

Ubiquitylation, and in particular cell cycle-regulated ubiquitylation, is often further regulated by additional post-translational modifications such as phosphorylation [61, 62, 63]. For instance, ubiquitylation and degradation of c-Mos after fertilization of *Xenopus* eggs is regulated by its phospho-status; addition of anti-Fizzy antibodies maintains high cyclin B/cdc2 and this prevents c-Mos dephosphorylation and its subsequent degradation, while anti-Fizzy antibodies had no effect on c-Mos dephosphorylation and destruction 15 minutes after activation, when cyclin B was already degraded [64]. Cyclin degradation is itself regulated by c-Mos, through the activation of MAP kinase which can prevents cyclin B-cdc2 kinase-triggered cyclin destruction [65]. By adding c-Mos protein to CSF-arrested egg extracts, the authors found that the cyclin degradation machinery was poised, but not inactivated, as release of the extract from the CSF block by addition of Ca2+-calmodulin-dependent protein kinase II allowed the degradation machinery to function normally [65].

Taken together, it is clear that work by the Kirschner [52, 45, 47, 55, 24, 58, 48, 57, 60], Hunt [19, 20, 66], and Lohka [50] labs in particular, alongside important contributions from a number of other researchers using the *Xenopus* extract system, has been formative in instructing our current understanding of cyclin destruction and cell cycle regulation of protein degradation.

## Ubiquitylation and DNA Replication

Protein degradation also plays a key role in regulation of DNA replication, and this has also been extensively characterized using *Xenopus* extracts. For instance, Geminin, an important negative regulator of DNA replication [67], which also has a distinct role in neural induction [68], was first identified by an unbiased screen set up to identify proteins destroyed specifically in mitosis [67]. Ub-mediated regulation of other DNA replication factors has also been studied by additional methods in *Xenopus* extracts. For instance, in the initiation of DNA replication, the requirement for Cdc34 as a Ub-conjugating enzyme to target proteins for degradation and drive entry into S-phase was established using methyl-Ub-supplemented *Xenopus* extracts [69]. DNA replication was inhibited by generally blocking Ub-mediated proteasomal degradation. It was also postulated that Cdc34 may regulate the cdk inhibitor Xic1s degradation and indeed in extracts with sperm nuclei added, Xic1 was efficiently degraded. It is possible to use antibodies to immunodeplete factors from egg extracts and then assay the affect on biochemical processes, and Xic1 was stable in Cdc34-depleted extracts [69]. This observation suggested a close link between Xic1 degradation and replication, in particular for formation of a prereplication complex requiring Cdk2, Cdc7 and Cdc45 before Xic1 could be degraded, while upon completion of DNA replication Xic1 is stabilized [70]. Findings that the Xic1 degradation rate correlated with the concentration of sperm nuclei in extract [69] were explained by the finding that chromatin and a nuclear environment were required for Xic1 degradation to occur [70]. Moreover, depletion of the chromatin-bound licensing factor Mcm7 from extracts inhibited both Xic1 degradation and DNA replication by preventing the formation of prereplication complexes on chromatin [70]. Therefore a variety of *Xenopus* extract depletion studies and degradation assays, previously used to demonstrate the cell cycle dependence of Ub-mediated protein degradation, have also been modified to highlight the relationship between ubiquitylation and DNA replication.

## Ub-mediated degradation of Neurogenin2; complex regulation in eggs and embryos

Ubiquitylation usually occurs on one or more lysine residues. However, other residues are nevertheless capable of forming bonds with Ub molecules and these can also target proteins for destruction. Non-canonical ubiquitylation describes the modification of protein substrates with Ub on residues other than the canonical lysine residue [71], which can occur onon the N-terminal amino group; the thiol group of cysteine residues; and the hydroxyl group of serine and threonine residues, all of which have the potential (and as shown in Tables 1, 2 and 3 of [13], the demonstrated activity) to react with and covalently link to Ub. Our own laboratory has investigated these non-canonical ubiquitylation events by looking in detail at the degradation of the proneural basic helix loop helix transcription factor Neurogenin2 (Ngn2, see Figure 2). This has proved to be surprisingly complex.

The original observations in *Xenopus laevis* interphase egg extract indicating that non-canonical ubiquitylation may be occurring where that Ngn2 protein, where all lysines had been mutated to alanines, could nevertheless be ubiquitylated and degraded efficiently. We found that Ub moieties could be directly added onto the N-terminus of Ngn2. Moreover, an N-terminally blocked lysine-less mutant of Ngn2 could be further ubiquitylated by bonds that were sensitive to reducing agents and showing pH-dependency, implicating Ub linkages to cysteines, serines and threonines ([32], as reviewed in [72, 34, 13]). Non-canonical ubiquitylation was of differing importance for Ngn2 protein stability in interphase versus mitosis; we saw that mutation of all the cysteines in Ngn2 had no effect on stability in interphase egg extracts, but was stabilizing in mitotic extracts [32]. Good evidence for non-canonical ubiquitylation of Ngn2 driving its destruction was also found in extracts from neurula-stage *Xenopus* embryos, indicating that this was not an egg-specific phenomenon. Indeed, similar non-canonical regulation was observed in mouse embryonal carcinoma cells, demonstrating that even unusual regulatory mechanisms first uncovered in *Xenopus* extracts are nevertheless active *in vivo* in frogs and in mammalian cells [25]. It is important to note, however, that we have since showed that the closely related proteins Ngn2 and Ngn3 show somewhat different control of ubiquitylation and destruction, and this demonstrates the possible need to consider these regulatory mechanisms on a protein by protein basis [73].

**Figure 2:**
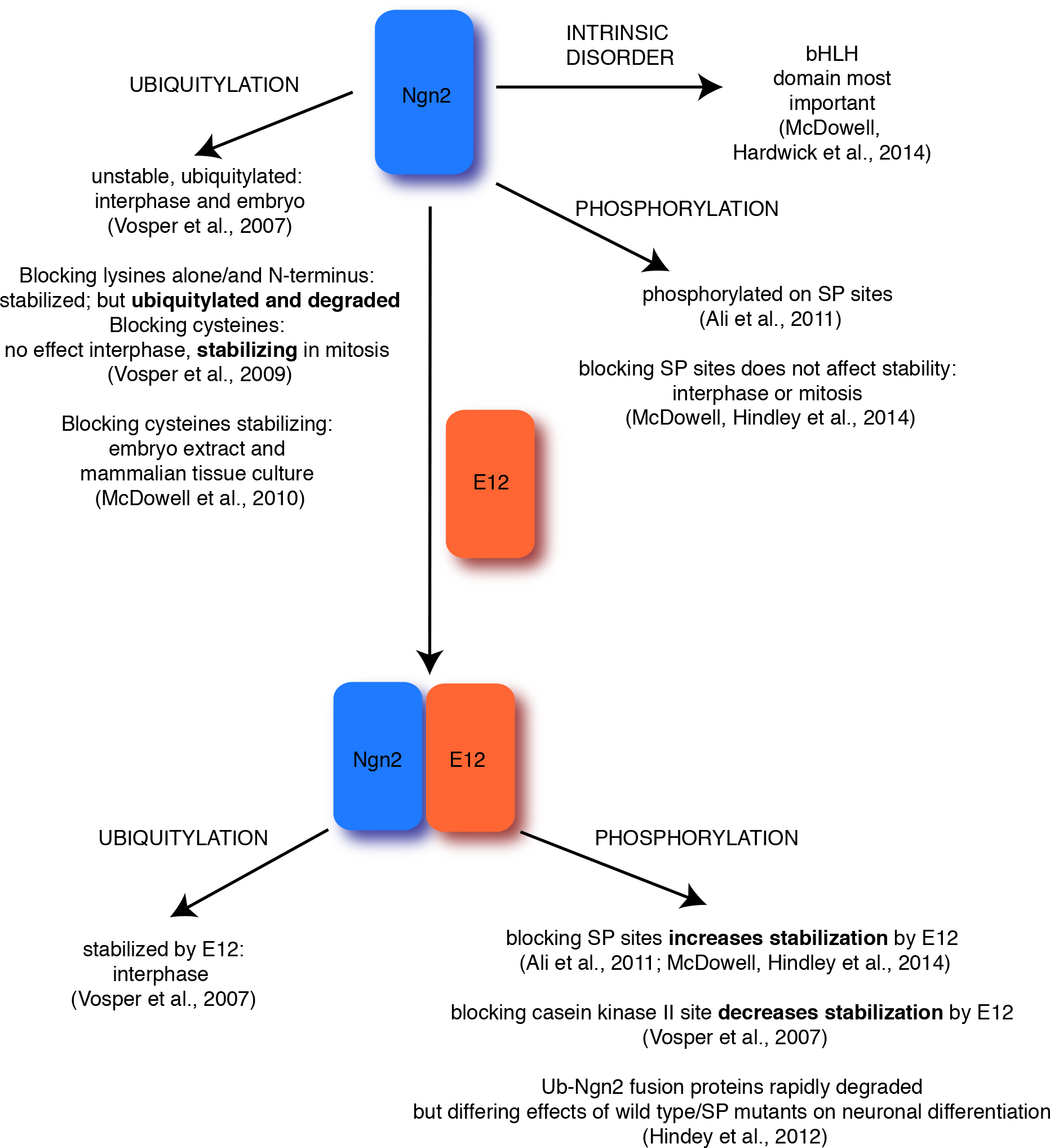
Regulation of Ngn2. Schematic illustrating points of regulation of ubiquitin-mediated degradation of Ngn2.

We have also undertaken a detailed investigation of the effects of cell cycle-regulated phosphorylation on Ngn2 protein activity [74, 75, 76] and it is now clear that the relationship between Ngn2 phosphorylation and protein stability is complex. The half-life of Ngn2 protein can be readily calculated by addition of radiolabelled *in vitro* translated Ngn2 protein to *Xenopus* extracts, removing samples at increasing times, and quantitating the amount of protein left after SDS PAGE and autoradiography ([77], Figure 1). Ngn2 is phosphorylated on multiple serine-proline motifs [74]. Mutation of these serines to alanines does not alter the half life of Ngn2 in interphase or mitotic extracts. However, Ngn2 protein is stabilized by addition of its heterodimeric E protein binding partner and this is regulated by the availability of these serines; the phosphomutant protein is more readily stabilized than the wild-type protein [74, 76]. This is in contrast to experiments looking at another potential casein kinase II phosphomutant of Ngn2, T118A, which was not as well protected from degradation as the wild type protein [77]. Moreover, fusion of Ub directly to the N-terminus of Ngn2 reduced its half-life dramatically, as well as significantly inhibiting its ability to drive neuronal differentiation, though the effect on neuronal differentiation of this fusion was less pronounced for the phosphomutant version of the protein [75]. These experiments, which use egg and embryo extracts in parallel with *in vivo* analyses, demonstrate clearly the versatility of *Xenopus* as a model system to study the functional consequences of ubiquitylation.

In addition to ubiquitylation (or sometimes even without ubiquitylation [78]), unfolding initiation sites are required to allow protein degradation by the UPS [79] and in particular proteins that are natively unfolded, lacking regular structure (intrinsically disordered or ID proteins [80]) may be particularly susceptible to Ub-mediated degradation [81, 13]. We investigated the role of ID in the degradation of Ngn2 using *Xenopus* extract systems [82], comparing Ngn2 and the related bHLH transcription factor NeuroD1, which is similarly structured but has very different degradation properties [77]. Both are ubiq-uitylated, but only Ngn2 is ubiquitylated on non-canonical sites [82], and it is degraded considerably faster that NeuroD. Using the extract system to compare the stability of a variety of chimeric proteins where we swapped N-, C- and bHLH domains between Ngn2 and NeuroD1, we found that despite the disorder in the N- and C-terminal domains likely in both proteins, the bHLH domain itself (which is still highly disordered [83] but has a much more highly conserved sequence between the two proteins [84]) was the region that appeared to most greatly influence chimeric protein activity and stability [82]. This analysis was only possible because of our ability to carry out a very large number of degradation assays using the *Xenopus* extract system. The other great advantage of *Xenopus* was also to have the *in vivo* system working in parallel with degradation assays to assess functional activity of the chimeric proteins. Together these allowed us to determine the relationship between structure, activity and stability. This relationship has proved to be remarkably complex ([74, 75, 82, 76], see Figure 2).

## Ubiquitylation in a developmental context

Most of the insights into ubiquitylation gained using *Xenopus* have come from biochemical analyses in extract systems as described above, and in general, most attention has been paid to degradation mechanisms found in the egg. However, many proteins are degraded at specific times in development, and there has been some work in both egg and embryo extracts that has addressed control of ubiquitylation and destruction of proteins only found expressed at later developmental stages (including Ngn2 as described above). Indeed, the great benefit of studying Ub-mediated protein degradation in *Xenopus* is the ability to put the biochemical information into a developmental context. For instance, Xic1 is targeted for degradation by the F-box protein Skp2, and its degradation has been studied in egg extracts as well as in embryos [85, 86]. The destruction of Skp2 itself by the APC/C has been studied in *Xenopus* extract by ubiquitylation and degradation assays [87] and its function analyzed in developing embryos [85].

One of the most interesting studies of Ub-mediated protein degradation in a developmental context is the study of *β*-catenin regulation in extracts from eggs and embryos, with implications for the study of Wnt signaling in embryogenesis [31]. Again, the strength of this approach lies in the ability to compare the effects of various treatments on *β*-catenin stability, as well as other components of the Wnt signaling pathway, with developmental defects that are observed, such as problems with axis formation (see Table 1, [31]).

## Concluding Remarks

Since the relatively recent discovery of Ub and acceptance of intracellular protein catabolism, degradation of proteins by the 26S proteasome via the Ub-proteasome system has become an area of intense study across biochemical, cell biology, developmental and clinical disciplines. From the earliest days of ubiquitylation research the “cell-in-a-test-tube” extract systems of the frog *Xenopus laevis* have catalyzed the discovery of components of this machinery. This utility is further strengthened because of the versatility of *Xenopus* as a developmental model, allowing investigation of the *in vivo* consequences of ubiquitylation along side *in vitro* assays more geared to biochemical characterisation. Overall, the availability of large quantities of extract from the eggs of a single frog in which ubiquitylation and protein degradation can occur, the speed with which experiments can be carried out, and the ability to compare *in vitro* findings with effects *in vivo* in a rapid and well-characterized developmental context, have made *Xenopus* a crucial model organism for studying ubiquitylation and protein degradation. Its high level of versatility means it will remain a highly valued system for continued studies of Ub-mediated degradation.

## Acknowledgements

Research in APs laboratory is supported by Medical Research Council grants MR/K018329/1 and MR/L021129/1, and core support from the Wellcome Trust and MRC to the Wellcome Trust — Medical Research Council Cambridge Stem Cell Institute. GM was supported by Michael Levin at Tufts University through grants from the G. Harold and Leila Y. Mathers Charitable Foundation and the Physical Science Oncology Center supported by Award Number U54CA143876 from the National Cancer Institute. GM is currently supported by a grant from Open Philanthropy and a residency funded by the Gordon and Betty Moore Foundation.

**Comments** Comments can be made by contacting the authors at.

